# Control of heat and oxidative stress adaptation by the DJ-1 paralogs in *Arabidopsis thaliana*

**DOI:** 10.1101/2024.06.05.597658

**Authors:** Priyanka Kataria, Naga Jyothi Pullagurla, Debabrata Laha, Patrick D’ Silva

## Abstract

Plant growth and development are highly regulated processes and are majorly controlled by various environmental factors, whose extreme exposures lead to chronic stress conditions promoting reactive oxygen species (ROS) and carbonyl species (RCS) production. ROS and RCS extensively damage cellular biomolecules and organelles, affecting plant’s viability and development. Emerging reports highlight that the multi-stress responding DJ-1 superfamily proteins are critical in attenuating cytotoxic effects associated with abiotic stress. The current report, validated in yeast and plant models, shows that *At*DJ-1C and *At*DJ-1E are robust antioxidants that scavenge ROS and improve survival under oxidative stress. Although they lack conventional glyoxalases and do not attenuate the glycation of proteins, *At*DJ-1C and *At*DJ-1E preserve the GSH pool and regulate redox homeostasis. Moreover, transcriptome profiling indicates that levels of *AtDJ-1C* and *AtDJ-1E* are rapidly established to counter heat and oxidative stress conditions. Notably, the knockdown of *AtDJ-1C* and *AtDJ-1E* promotes detrimental alterations such as reduced chlorophyll retention, impaired root morphogenesis, and induced sensitivity to heat stress due to ROS elevation. Contrastingly, overexpression of *At*DJ-1C and *At*DJ-1E improved plant height and rosette formation under physiological conditions. In conclusion, our study unravels the pivotal functions of *Arabidopsis thaliana* DJ-1C and DJ-1E in governing plant health and survival under heat and oxidative stress conditions.

## Introduction

Various environmental challenges, including extreme temperatures, salinity, drought, and heavy metal toxicity, affect plant growth, development and crop productivity. Prolonged exposure to these stresses leads to inhibition in seed germination, reduced stomatal conductance, decrease in absorption and delayed maturation (Sachdev et al., 2021). At the molecular level, these environmental stresses augment the generation of Reactive Oxygen Species (ROS) and Reactive Carbonyl **S**pecies (RCS). However, under physiological conditions, they play an important role in signalling, cell differentiation, and development as the balance is maintained between the generation and removal of these reactive species (Hoque et al., 2012; Mittler et al., 2022). Their increased production during stressful conditions causes damage to nucleic acids (DNA and RNA), oxidation of proteins and peroxidation of lipids, leading to alteration in photosynthetic apparatus, photorespiration and mitochondrial respiration (Tripathi et al., 2023). Therefore, governing their cellular concentrations and preventing aberrant damage are the foremost requirements in the cellular milieu. Plants have two major antioxidant systems: enzymatic (including superoxide dismutase, catalase, glutathione reductase, etc.) and non-enzymatic (including glutathione, carotenoids, etc.) to minimize the detrimental effect of ROS on cellular macromolecules. On the contrary, they have evolved with a dedicated glyoxalase system involving glyoxalase I and II enzymes, which detoxifies RCS (Methylglyoxal {MG} and Glyoxal {GO}) by utilizing glutathione (GSH) as a co-factor, which becomes limiting during oxidative stress. This limitation is overcome by the GSH-independent glyoxalase III system, which involves ThiJ/DJ-1/PfpI superfamily member proteins.

DJ-1 superfamily member proteins are present across the phylogeny from prokaryotes to eukaryotes with varying numbers of paralogs. Apart from glyoxalase activity, they exhibit various functions, including mitochondrial regulation, chaperonin activity, deglycase activity, transcriptional regulation and antioxidant (Zhou et al., 2006; Tsai et al., 2015; Bankapalli et al., 2020; Richarme et al., 2018). Human DJ-1, which is encoded by the PARK7 gene, is the most studied protein of the DJ-1 superfamily since its mutations cause familial cases of Parkinson’s disease. It is a multifunctional protein that protects neurons from oxidative stress, neurotoxins, and misfolded proteins by regulating mitochondrial health (Junn et al., 2009). In *S. cerevisiae*, four paralogs (Hsp31, Hsp32, Hsp33, Hsp34) are present, which are substrate-specific glyoxalases, stress-induced chaperones, and their absence induces oxidative and carbonyl stress in cells (Bankapalli et al., 2015; Tsai et al., 2015; Susarla et al., 2023). Recently, DJ-1 proteins have also been referred to as deglycase, where they repair methylglyoxal and glyoxal glycated nucleic acids and proteins (Richarme et al., 2015; Sharma et al., 2019; Zheng et al., 2019; Susarla et al., 2023; Richarme et al., 2018).

In plants, the physiological role of DJ-1 members remained largely unexplored, with few studies indicating their role in various environmental stress conditions. Interestingly, a comprehensive survey reports that the plant kingdom possesses a large number of *DJ-1* isoforms in comparison to other organisms (Ghosh et al., 2016). Moreover, they have evolved with two DJ-1 domains containing two catalytic sites in a single polypeptide backbone. Intriguingly, 27 out of 217 sequences were found to have a single domain, which specifies the preference towards having a double domain. Comparative transcriptome profiling suggests variable expression patterns of *DJ-1* genes in different plant tissues, indicating their tissue-specific functions at different stages of development (Ghosh, 2017a; Islam and Ghosh, 2018; Prasad et al., 2022). Apart from glyoxalase activity, plant DJ-1 members of sugarcane (*Erianthus arundinaceous),* tomato *(Solanum lycopersicum)*, date palm (*Phoenix dactylifera*) and rice (*Oryza sativa*) are also involved in stress tolerance towards abiotic stress conditions such as drought, salinity and osmotic stress (Mohanan et al., 2020a; Jana et al., 2021; Gambhir et al., 2023b; Rathore et al., 2024). Overexpression of Hsp31 from *S. Cerevisiae* augments tolerance against abiotic and biotic stresses in tobacco (*Nicotiana tabacum*), suggesting their conserved functions across the kingdom of life (Melvin et al., 2017). *Arabidopsis thaliana* genome encodes six DJ-1 paralogs [DJ-1A to DJ-1F] with diverse physiological functions. The expression level of *AtDJ-1A* is upregulated in response to various exogenous cues, and consequently, it stimulates the activity of CSD1 and SOD1 (Xu et al., 2010a). On the other hand, *At*DJ-1B was reported as a bifunctional protein having glyoxalase and holdase activity, where its glyoxalase activity was lost upon oxidative stress (Lewandowska et al., 2019). Moreover, the recent finding on *At*DJ-1D highlights that it is a multifunctional protein that has glyoxalase, antioxidants, deglycase, and chaperone activity. Furthermore, overexpression of *At*DJ-1D contributes to the elevated tolerance towards carbonyl and oxidative stress, which was validated in both yeast and plant model species (Choi et al., 2014; Prasad et al., 2022; Gambhir et al., 2023a)

In our current study, we show that *Arabidopsis* DJ-1C and DJ-1E lack conventional glyoxalase activity and do not alter the glycation status of the macromolecules. Their knockdown resulted in poor plant development and enhanced ROS levels during heat and oxidative stress. We also discovered that they scavenge ROS and promote redox homeostasis by regulating the global glutathione levels. Together, these diverse properties of *At*DJ-1C and *At*DJ-1E play a pivotal role in providing superior growth with longer root lengths and higher chlorophyll retention under the most prevalent abiotic stress conditions, such as thermal and oxidative stress. Besides providing robust cellular protection under abiotic stress, their overexpression leads to morphological alterations such as enhanced rosette formation and increased plant height.

## Results

### *Arabidopsis* DJ-1C and DJ-1E do not detoxify reactive carbonyl species

Plants have evolved multiple homologs of DJ-1 with distinct physiological functions, enabling a robust stress adaptation. *Arabidopsis* possesses six DJ-1 paralogs (DJ-1A to DJ-1F) [Supplementary Fig. 1A], which are conserved and contain two distinct DJ-1 domains within a single polypeptide backbone, displaying a unique protein morphology. Our previous study reported that *Arabidopsis* DJ-1D is a novel enzyme that scavenges MG, GO, and ROS, further regulating plant physiology and health (Prasad et al., 2022). To investigate the physiological role of *Arabidopsis* DJ-1 paralogs, *At*DJ-1C and *At*DJ-1E, first, we studied the consequences of heterologous expression of these paralogs in the budding yeast *Saccharomyces cerevisiae*. Both of the paralogs were cloned into yeast vector (pRS415) and were expressed in the Δ*glo1* yeast knockout strain, as glyoxalase I is extensively involved in detoxifying carbonyls in yeast. The respective yeast transformants were subjected to phenotypic analysis under MG and GO stress. As expected, the Δ*glo1* mutant displayed severe growth sensitivity when exposed to both stress conditions. Strikingly, overexpression of *AtDJ-1C* and *AtDJ-1E* did not restore the growth defect of Δ*glo1* yeast mutant under both MG and GO stress. Notably, ectopic expression of *AtDJ-1D* which is a known glyoxalase, fully rescues the Δ*glo1* yeast-associated growth defects [Fig.1 A]. We also analysed the steady-state protein levels of DJ-1 paralogs of Δ*glo1* yeast transformants by using an anti-HA antibody [Supplementary Fig. 1B]. Since MG and GO are potent glycating agents, the proteome glycation profile was analyzed using anti-MAGE and anti-CML antibodies, which probe for MG and GO glycated proteins. Cells from the mid-log phase were treated with either MG or GO and processed for western analysis. As observed earlier, the Δ*glo1* strain showed enhanced protein glycation in comparison to the Col-0 (wild-type) and *DJ-1D* overexpression strains [Fig.1 B and C]. However, the ectopic expression of *At*DJ-1C and *At*DJ-1E did not restore the glycation levels of the GLO1-deficient yeast strain, suggesting that *Arabidopsis* DJ-1C and DJ-1E do not attenuate MG and GO-induced glycation in *Saccharomyces cerevisiae* [Fig.1 B and C].

**Figure. 1:**
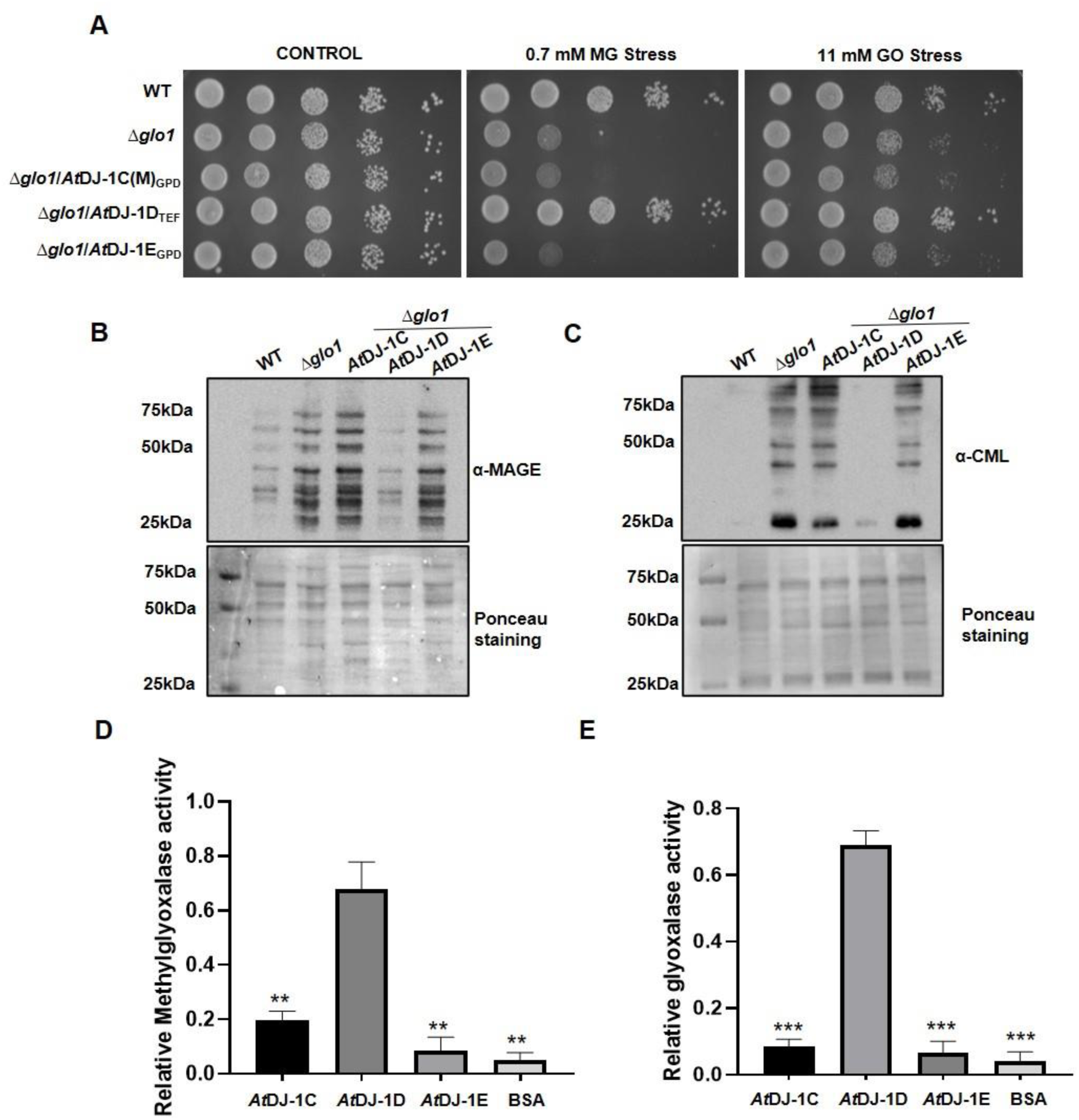
*At*DJ-1C and *At*DJ-1E fail to scavenge methylglyoxal and glyoxal. **(A) Overexpression of *At*DJ-1C and *At*DJ-1E does not provide protection from carbonyl stress.** Transformants expressing *At*DJ-1C and *At*DJ-1E in Δ*glo* strain of *Saccharomyces cerevisiae* were grown till mid-log phase and spotted in 10-fold serial dilutions onto SD Leu^-^ plates with or without 0.7 mM methylglyoxal or 11 mM glyoxal. Images were captured after 36 h. **(B and C) *At*DJ-1C and *At*DJ-1E do not attenuate protein glycation in yeast**. The respective strains treated with 2 mM methylglyoxal and 18 mM glyoxal for 6 h were subjected to immunoblotting using antibodies **(B)** anti-MAGE antibody for MG-glycated proteins and **(C)** anti-CML antibody for GO-glycated proteins. Ponceau stain was used as loading control. **(D and E) *At*DJ-1C and *At*DJ-1E do not possess glyoxalase activity.** Purified proteins, *At*DJ-1C, *At*DJ-1D, and *At*DJ-1E, were subjected to *in vitro* methylglyoxalase and glyoxalase activity assay with *At*DJ-1D as positive control and BSA as negative control. Data from three representative experiments are (mean SE; one-way ANOVA: *, *P* ≤ 0.05; **, *P* ≤ 0.01; ***, *P* ≤ 0.001; ****, *P* ≤ 0.0001; ns, not significant).

To further explore the enzymatic properties of these proteins, we purified 6X-histidine-tagged *At*DJ-1C (without N-terminal CTP signal sequence), *At*DJ-1D, and *At*DJ-1E using Ni^2+^ NTA affinity chromatography from *E. coli* [Supplementary Fig. 1C]. Using these recombinant proteins, we performed *in vitro* glyoxalase assay, and observed that *At*DJ-1C and *At*DJ-1E could not scavenge MG and GO compared to *At*DJ-1D, which is a robust scavenger of carbonyls (Prasad et al., 2022) [Fig.1 D and E]. BSA was used as the negative control. Together, these findings suggest that *At*DJ-1C and *At*DJ-1E fail to detoxify carbonyls and do not regulate protein glycation.

### *Arabidopsis* DJ-1 paralogs contribute to oxidative stress resistance in *Saccharomyces cerevisiae*

Reactive oxygen species are inevitably generated during various metabolic pathways in multiple organelles, like mitochondria, chloroplast, and peroxisomes; their basal levels are responsible for the signalling cascade(Waszczak et al., 2018). The cellular ROS is further elevated during exposures to salinity, drought, and environmental changes, leading to oxidative damage. DJ-1 family members from yeast, plants, and humans are known to regulate ROS homeostasis (Taira et al., 2004; Xu et al., 2010b; Bankapalli et al., 2015). Since *Arabidopsis* DJ-1C and DJ-1E do not possess canonical methylglyoxalase and glyoxalase activity [Fig. 1D and E], we then determined their antioxidant property in *S. cerevisiae* by heterologously expressing *AtDJ-1C*, *AtDJ-1E*, and *AtDJ-1D* in a ROS-sensitive strain, Δ*hsp31*. Upon 1 mM H_2_O_2_ stress, *At*DJ-1C and *At*DJ-1E rescued the growth phenotype of Δ*hsp31*, comparable to *At*DJ-1D, an established antioxidant, and WT [Fig. 2A].

**Figure. 2:**
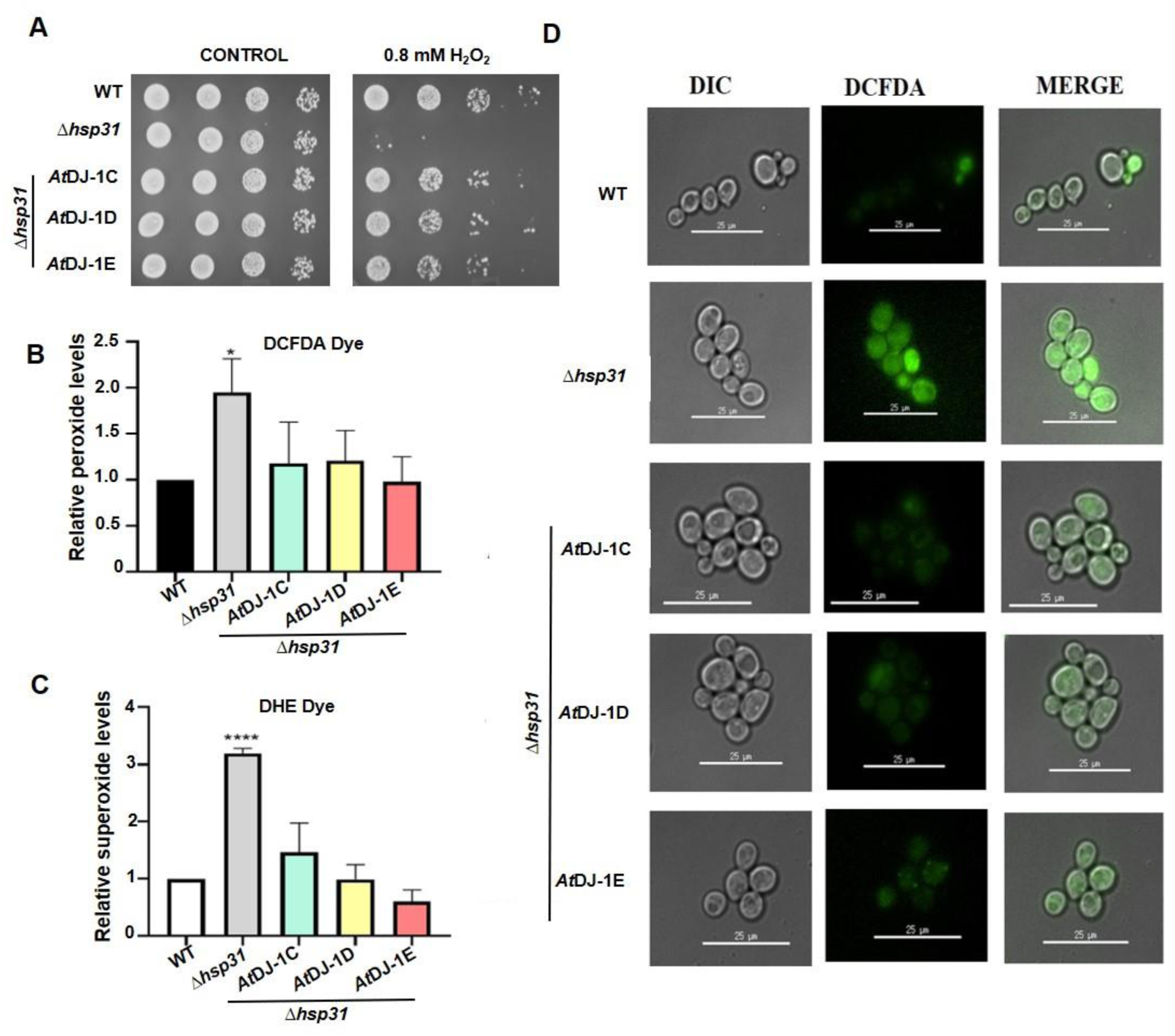
Ectopic expression of *At*DJ-1C and *At*DJ-1E provides tolerance to oxidative stress in *Saccharomyces cerevisiae*. **(A) *At*DJ-1 paralogs provide oxidative stress resistance.** Spot assay showing DJ-1 paralogs rescues growth defect of Δ*hsp31* under oxidative stress. WT (BY4741), Δ*hsp31* and Δ*hsp31* overexpressing DJ-1 paralogs strains were treated with water (*right panel*) or 0.8 mM H2O2 (*left panel*) and spotted in 10-fold serial dilutions onto SD leu^-^ media. Images were captured after 48 h. (**B, C and D) Estimation of cytosolic ROS levels**. An equivalent amount of cells from each strain were pelleted and exposed with 1 mM H2O2. Treated strains were stained with **(B)** DCFDA and **(C)** DHE dye, followed by flow cytometric analysis. Data from three representative experiments are shown (mean SE; one-way ANOVA: *, *P* ≤ 0.05; **, *P* ≤ 0.01; ***, *P* ≤ 0.001; ****, *P* ≤ 0.0001; ns, not significant). **(D)** Fluorescence microscopy showing DCFDA dye staining (green fluorescence) in WT (BY4741), Δ*hsp31* and Δ*hsp31* overexpressing DJ-1 paralogs strains upon 1 mM H2O2 stress by using Delta Vision Elite microscope

To further support the findings, we performed flow cytometric analysis with H_2_DCFDA and DHE dyes, which measure peroxide and superoxide levels, respectively. As reported, the absence of Hsp31 aggravated ROS levels compared to WT (Bankapalli et al., 2015). Interestingly, *At*DJ-1C and *At*DJ-1E significantly repressed the accretion of peroxide and superoxide levels, similar to *At*DJ-1D [Fig. 2B and C]. Next, fluorescence microscopy was performed to visualize the intensity of H_2_DCFDA dye between the strains. In agreement with flow cytometric data, Δ*hsp31* displayed higher ROS levels, indicated by the intense green fluorescence, while overexpression of DJ-1 paralogs attenuated the amounts of both peroxide and superoxide [Fig. 2D]. These results cumulatively suggest that *Arabidopsis At*DJ-1C and *At*DJ-1E reduce global cellular ROS and provide oxidative stress resistance in yeast.

### Knockdown lines of *At*DJ-1C and *At*DJ-1E are sensitive to high-temperature and oxidative stress

Abiotic stresses such as extreme temperatures (heat or cold), drought, and salinity are often the primary inducers of cellular damage and are also responsible for variation in the transcriptome in plants (Kreps et al., 2002; Vashisth et al., 2018). Such essential reprogramming is a feature of stress adaptation. We, therefore, examined the expression profile of *AtDJ-1C* and *AtDJ-1E* in diverse environmental challenges including heat, cold, salinity, carbonyl stress, osmotic and oxidative stress. RNA was extracted from seedlings treated without or with the above-mentioned conditions and subjected to qRT-PCR, *AtDJ-1C* and *AtDJ-1E* showed a substantial four to eight-fold upregulation in transcript levels in response to oxidative and heat stress [Fig. 3A and B]. However, only *AtDJ-1E* was upregulated during cold stress in comparison to *AtDJ-1C* [Supplementary Fig. 2A], while both genes were found to be downregulated in the presence of carbonyl and osmotic stress [Fig. 3C and Supplementary Fig. 2B]. On the other hand, only *AtDJ-1C* showed downregulation upon salt stress in comparison to *AtDJ-1E*, whose levels were not affected significantly [Supplementary Fig. 2C].

**Figure 3.**
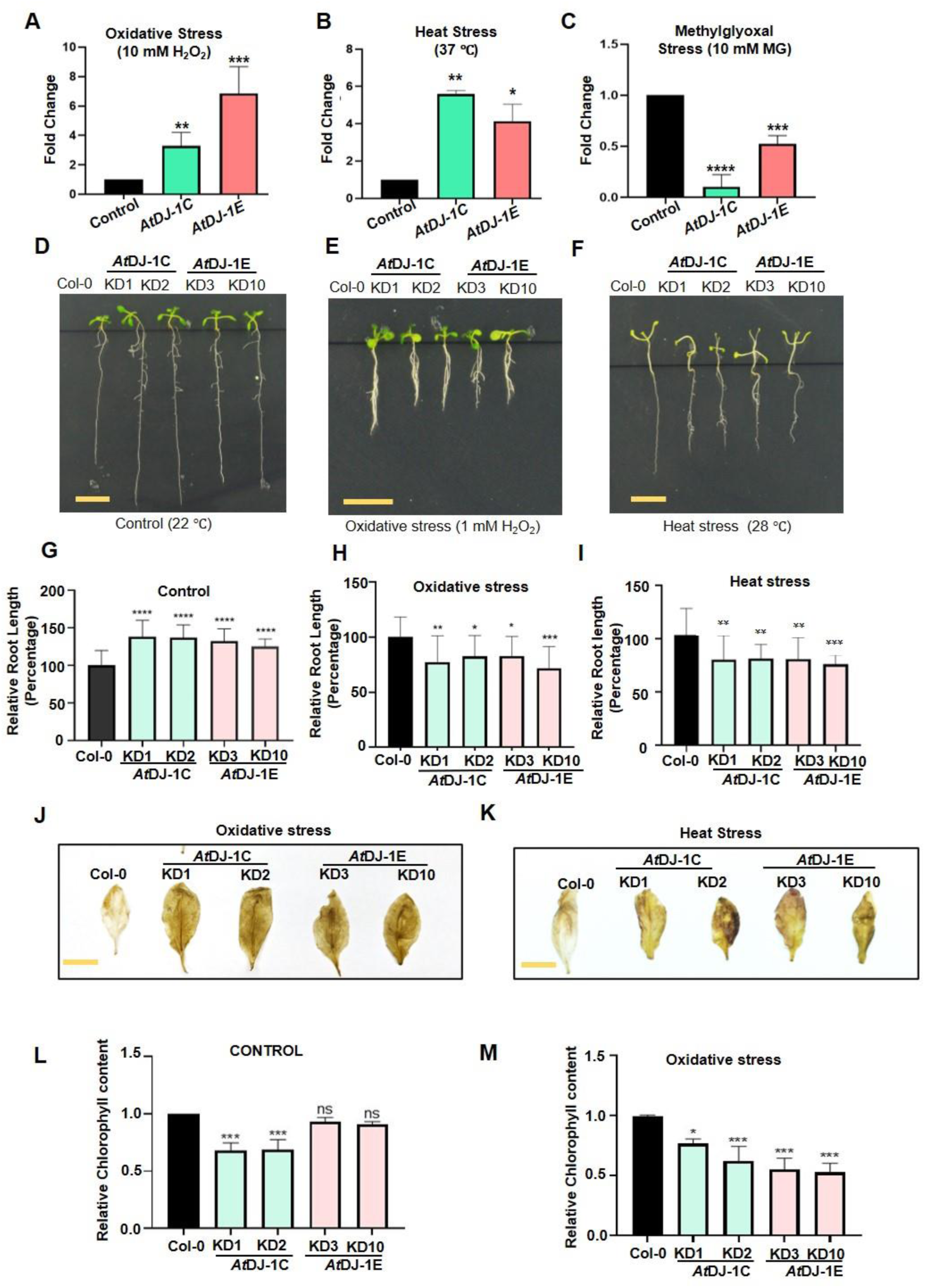
*Arabidopsis* plants with compromised expression of *AtDJ-1C* and *AtDJ-1E*, exhibit altered root development during oxidative stress and heat treatment. **(A-C) Differential expression of *DJ-1C* and *DJ-1E* during various abiotic stresses.** Quantitative RT-PCR (qRT-PCR) analysis of *AtDJ-1C* and *AtDJ-1E* in Col-0 under **(A)** oxidative stress, **(B)** heat stress and **(C)** methylglyoxal stress. Seven-day-old seedlings were exposed to the above-mentioned stresses for 3 h and were harvested for qRT-PCR analysis. TUBULIN served as a control. Data from three biological replicates are shown (mean SE; one-way ANOVA: *, *P* ≤ 0.05; **, *P* ≤ 0.01; ***, *P* ≤ 0.001; ****, *P* ≤ 0.0001; ns, not significant). **(D-I) *At*DJ-1C and *At*DJ-1E regulate root morphogenesis.** 7-day-old seedlings of the designated genotypes were grown at **(D)** control condition or **(E)** exposed to 28°C or **(F)** treated with 0.6 mM H2O2 and allowed to grow for another 7 days followed by imaging. Representative images of 14-day-old seedlings of the indicated genotypes are presented. Scale bar = 1 cm. **(G-I) Graphs representing quantification of root lengths.** Root lengths of Col-0, independent knockdown lines of *AtDJ-1C* and *AtDJ-1E* under control conditions, heat stress and oxidative stress were measured using ImageJ. Data from 15-18 seedlings were taken and shown (mean SE; one-way ANOVA: *, *P* ≤ 0.05; **, *P* ≤ 0.01; ***, *P* ≤ 0.001; ****, *P* ≤ 0.0001; ns, not significant). **(J-K) Knockdown lines of *At*DJ-1C and *At*DJ-1E show reduced chlorophyll retention.** The leaves were collected from 4-week-old plants, and chlorophyll content was estimated by acetone extraction method under **(J)** control conditions and **(K)** oxidative stress (5 mM H2O2 for 24 h). Data from three representative experiments are shown (mean SE; one-way ANOVA: *, *P* ≤ 0.05; **, *P* ≤ 0.01; ***, *P* ≤ 0.001; ****, *P* ≤ 0.0001; ns, not significant). **(L-M) *At*DJ-1C and *At*DJ-1E absence contribute to peroxide accumulation during abiotic stress.** Leaves of *At*DJ-1C and *At*DJ-1E knockdown lines were exposed to **(L)** oxidative stress (5 mM H2O2 for 3 h) and **(M)** thermal stress (12 h at 30°C). Post-treatment, leaves were stained with 3,3′-diaminobenzidine (DAB) for 5 h, and images were taken after chlorophyll bleaching.

Since transcript levels of both *At*DJ-1C and *At*DJ-1E were upregulated during oxidative and heat stresses, we aimed to characterize *Arabidopsis* lines defective in *At*DJ-1C and *At*DJ-1E. Unfortunately, we were unable to isolate homozygous T-DNA knockout lines for both *AtDJ-1C and AtDJ-1E*; this is in agreement with the previously published report(Lin et al., 2011a). Therefore, we decided to work with heterozygous lines with reduced transcript of the respective paralogs [Supplementary Fig. 2D-F]. To assess the consequences of compromised expression of *At*DJ-1C and *At*DJ-1E, we examined the primary root development of *AtDJ-1C* and *AtDJ-1E* knockdown lines along with Col-0 (wild-type) plants. Intriguingly, under controlled conditions (22 °C), *AtDJ-1C* and *AtDJ-1E* knockdown lines displayed longer primary root lengths than Col-0 [Fig. 3D and G]. However, upon oxidative and heat stress, root elongation in independent knockdown lines was repressed when compared with Col-0 plants, suggesting that the activity of *At*DJ-1C and *At*DJ-1E is critical to regulate root development during oxidative and heat stresses [Fig. 3E, F, H and I].

Harsh environmental conditions affect the functionality of various plant parts, including leaves that house chloroplast, mitochondria and essential organelles. The damage to the leaves contributes to accelerated ROS production, which could be detected using DAB (3,3′-diaminobenzidine) staining (Daudi et al., 2012). Upon exposure of plants to heat and oxidative stress, we detected significant accumulation of peroxide levels in the independent *AtDJ-1C* and *AtDJ-1E* knockdown lines, indicated by the browning of leaves, as compared to Col-0 [Fig. 3 J and K]. Furthermore, we determined chlorophyll content in mutant lines, which is a hallmark of plant growth and survival as stress conditions primarily target photosynthetic apparatus and impair photosynthetic capacity (Hu et al., 2020). In control conditions, chlorophyll content in *AtDJ-1C* knockdown lines was markedly lower than Col-0 and *AtDJ-1E* knockdown lines [Fig. 3L]. Intriguingly, independent knockdown lines of *AtDJ-1C* and *AtDJ-1E* displayed substantially attenuated chlorophyll accumulation upon oxidative stress when compared with the Col-0 plants [Fig. 3M]. To summarize, *At*DJ-1C and *At*DJ-1E are critical antioxidants that regulate redox homeostasis in both yeast and plant models and further regulate chlorophyll content and plant physiology under heat and oxidative stress.

### Overexpression of DJ-1C and DJ-1E provides tolerance towards heat and oxidative stress

To further investigate the role of *Arabidopsis* DJ-1 paralogs in heat and oxidative stress, we generated two independent *AtDJ-1C* and *AtDJ-1E* overexpression lines. Both genes were expressed under the control of a constitutive CaMV 35S promoter having an N-terminal translational fusion with GFP. Stable *AtDJ-1C* and *At*DJ-1E overexpression lines were confirmed by PCR-based genotyping [Supplementary Fig. 3A and B]. Under control growth conditions, independent overexpression lines of *AtDJ-1C* and *AtDJ-1E* showed a superior growth phenotype, including enlarged rosette formation and increased plant height, compared to Col-0 plants [Supplementary Fig. 3C and Fig. 4A]. Notably, the overexpression lines did not show any substantial difference in primary root elongation (22 °C) [Fig. 4B and E]. Intriguingly, *AtDJ-1C* and *AtDJ-1E* overexpression lines exhibited longer primary root lengths than Col-0 when exposed to heat and oxidative stress, indicating their involvement in root development and maintenance during abiotic stress [Fig. 4C, D, F, and G].

**Figure 4.**
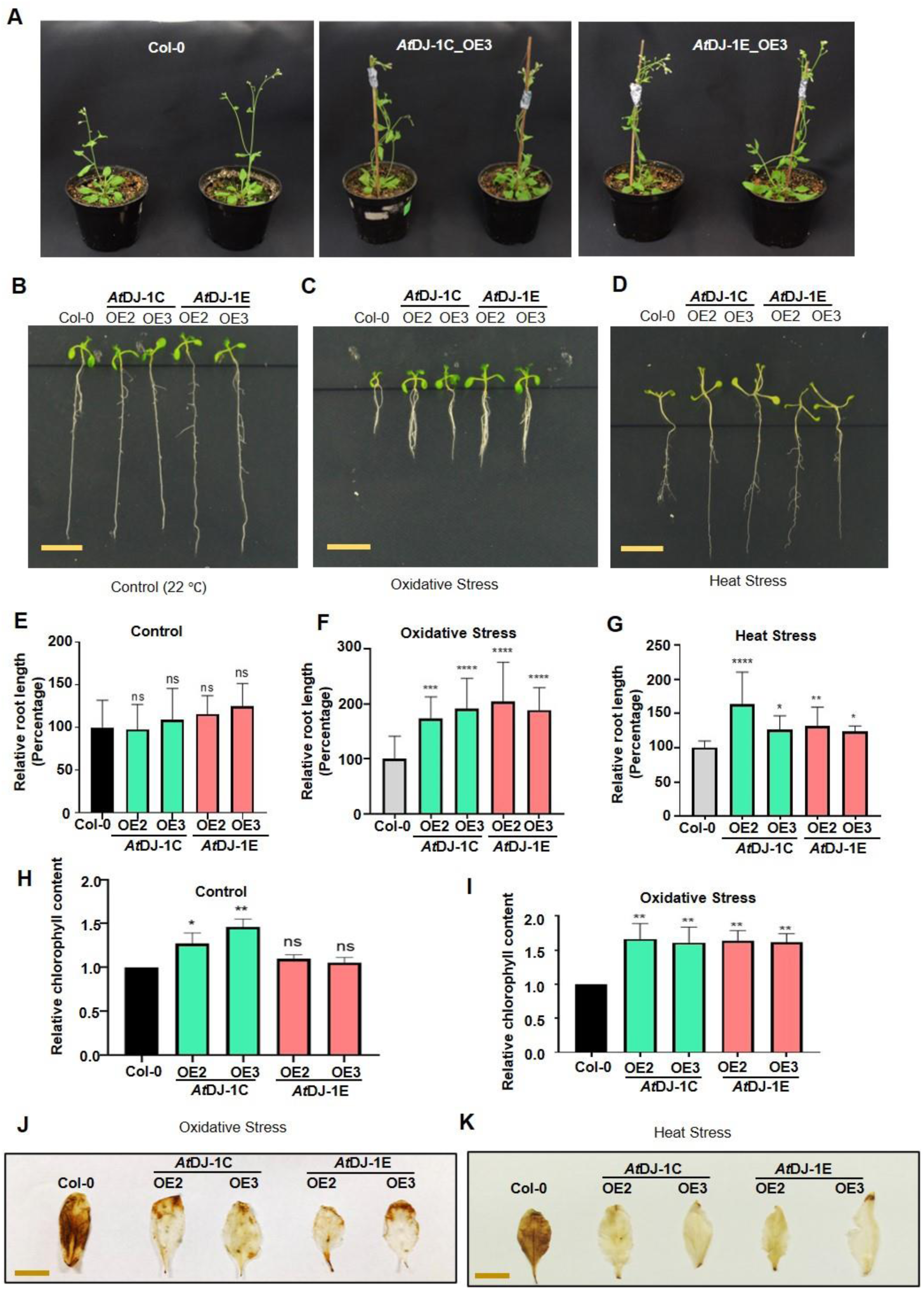
*At*DJ-1C and *At*DJ-1E overexpression lines display tolerance to oxidative and thermal stress in plants. **(A) *Arabidopsis* DJ-1 overexpression lines exhibit superior plant growth.** Representative pictures of 6-week-old wild-type (Col-0) and transgenic lines expressing GFP tagged DJ-1C and DJ-1E. **(B-D) *At*DJ-1C and *At*DJ-1E regulate root morphogenesis under abiotic stress.** Representative images of 2-week-old wild-type (Col-0) and transgenic lines expressing GFP tagged DJ-1C and DJ-1E grown in the presence or absence of oxidative and heat stress. 7 days old seedlings were transferred to solidified half-strength MS media supplemented with 1% (w/v) sucrose and 1 mM H2O2. The growth of root length was evaluated under (**B**) control, **(C)** oxidative stress and **(D)** heat stress using ImageJ software. **(E-G) Graphical representation of root lengths.** The percentage of changes in primary root length altered by provided oxidative and heat stress was determined for the Col-0, *At*DJ-1C and *At*DJ-1E independent overexpression lines. The relative root elongation of the designated genotypes under **(E)** control condition **(F)** oxidative and **(G)** heat stress. Data from 15-18 seedlings were taken and shown (mean SE; one-way ANOVA: *, *P* ≤ 0.05; **, *P* ≤ 0.01; ***, *P* ≤ 0.001; ****, *P* ≤ 0.0001; ns, not significant). **(H-I) *At*DJ-1C and *At*DJ-1E overexpression lines display higher chlorophyll retention.** Leaves were collected from 4-week-old plants, and chlorophyll content was estimated by acetone extraction method under **(H)** control conditions and **(I)** oxidative stress (5 mM H2O2). Data from three representative experiments are shown (mean SE; one-way ANOVA: *, *P* ≤ 0.05; **, *P* ≤ 0.01; ***, *P* ≤ 0.001; ****, *P* ≤ 0.0001; ns, not significant). **(J-K) Overexpression of *At*DJ-1C and *At*DJ-1E results in lower peroxide accumulation.** Leaves of *At*DJ-1C and *At*DJ-1E overexpression lines were exposed to **(J)** oxidative stress (5 mM H2O2 for 12 h) and **(K)** thermal stress (24 h at 30°C). Post-treatment, leaves were stained with 3,3′-diaminobenzidine (DAB) for 5 h, and images were taken after chlorophyll bleaching.

Furthermore, chlorophyll accumulation was also quantitated in the respective lines, and higher chlorophyll content was observed in the *AtDJ-1C* overexpression lines than in Col-0 and *AtDJ-1E* overexpression lines under control conditions [Fig. 4H]. Strikingly, both the overexpression lines showed enhanced chlorophyll retention than Col-0 during oxidative stress [Fig. 4I]. We also examined the distribution of peroxide levels in leaf tissues of *AtDJ-1C* and *AtDJ-1E* overexpression lines. Upon DAB staining, no significant difference was observed between Col-0 and overexpression lines under control conditions [Supplementary Fig. 3D]. However, exposure to heat and oxidative stress resulted in lower ROS levels in *AtDJ-1C* and *AtDJ-1E* overexpression lines when compared with Col-0 plants [Fig. 4J and K]. Together, these results suggest that *Arabidopsis* DJ-1C and DJ-1E are pivotal in governing plant health, growth, and physiology by attenuating ROS and suppressing thermal-associated damage.

### DJ-1 paralogs maintain glutathione levels in plants during oxidative stress

Recent studies suggest the role of DJ-1 homologs in glutathione (GSH) homeostasis, a thiol-containing tripeptide that is a major non-enzymatic antioxidant ubiquitously present in various sub-cellular compartments to scavenge ROS(Gambhir et al., 2023b). Moreover, the lack of DJ-1 members in *S. cerevisiae* was reported to abrogate the GSH pool (Bankapalli et al., 2015). Therefore, to understand the molecular mechanism by which Arabidopsis DJ-1C and DJ-1E offer ROS protection, we measured GSH levels in the presence of *Arabidopsis* DJ-1 paralogs in the *S. cerevisiae* Δ*hsp31* strain defective in yeast DJ-1 homolog. The membrane-permeable monochlorobimane (MCB) dye was utilized to monitor cellular levels of GSH. In agreement with the previous report (Bankapalli et al., 2015), our microscopic study reveals that the lack of Hsp31 leads to a significant reduction in the GSH pool, indicated by poor MCB signal [Fig. 5A]. On the contrary, overexpression of DJ-1C and DJ-1E restored the glutathione levels of Δ*hsp31* [Fig. 5A]. Furthermore, we performed flow cytometric analysis with MCB dye and observed that overexpression of *At*DJ-1C and *At*DJ-1E substantially restored the GSH levels [Fig. 5B].

**Figure 5.**
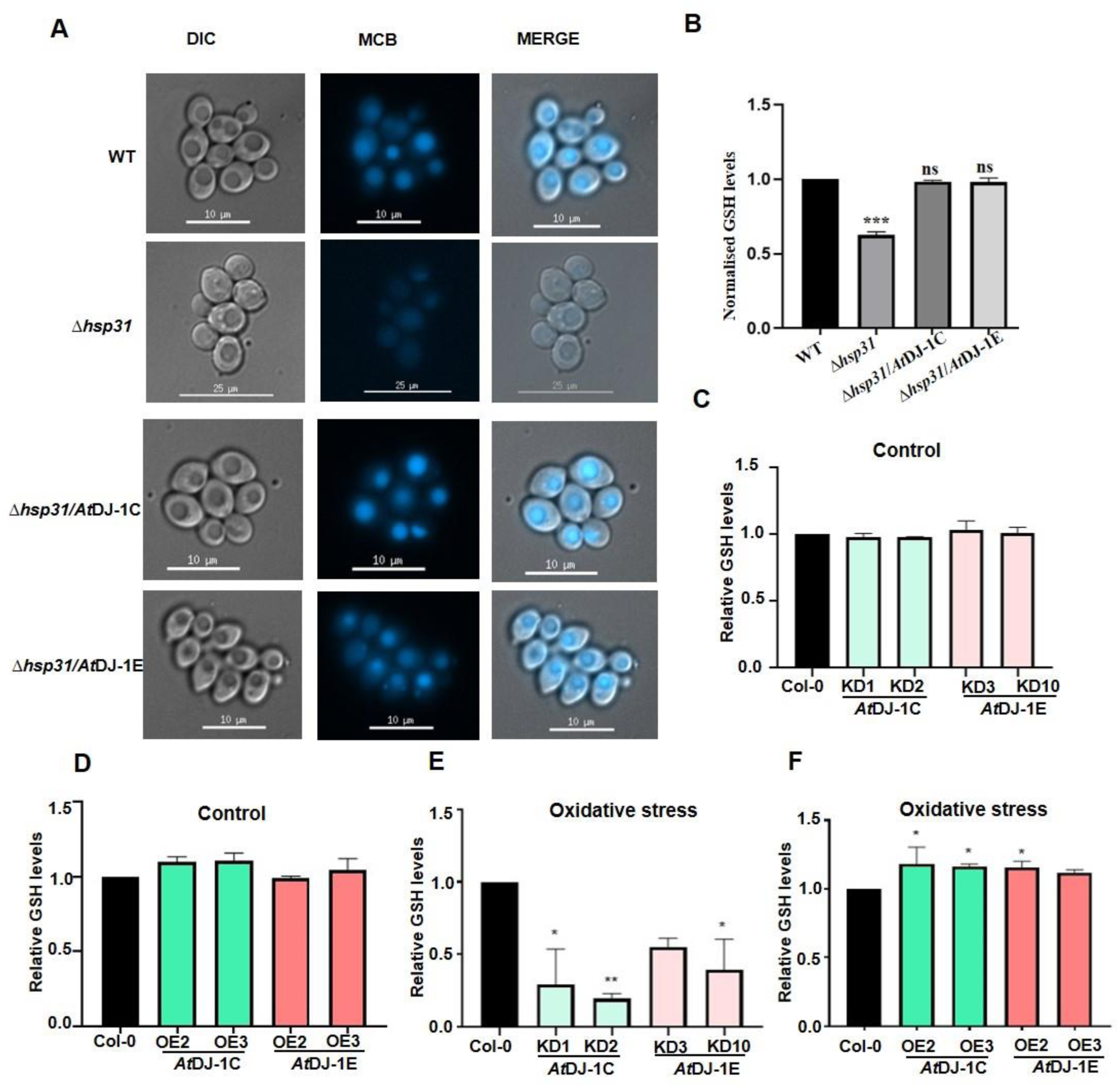
*At*DJ-1C and *At*DJ-1E regulate glutathione levels *in vivo*. **(A-B) Ectopic expression of *At*DJ-1C and *At*DJ-1E restores glutathione levels in *S. cerevisiae*.** WT (BY4741), Δ*hsp31* and Δ*hsp31* overexpressing DJ-1 paralogs strains grown till the mid-exponential phase were exposed to 1 mM H2O2 and stained with monochlorobimane (MCB) dye and visualized in **(A)** Fluorescence microscopy (blue fluorescence) and quantified the GSH levels in **(B)** flow cytometric analysis. Data from three representative experiments are shown (mean SE; one-way ANOVA: *, *P* ≤ 0.05; **, *P* ≤ 0.01; ***, *P* ≤ 0.001; ****, *P* ≤ 0.0001; ns, not significant). **(C-F) DJ-1 paralogs maintain glutathione levels in *Arabidopsis* during oxidative stress.** 4-week-old plants were subjected to glutathione estimation under control conditions in (C) *At*DJ-1C and *At*DJ-1E knockdown lines and **(D)** overexpression lines using a GSH estimation kit. 4-week-old plant leaves were treated with 5 mM H2O2 for 12 h to determine the GSH levels in **(E)** knockdown lines and **(F)** overexpression lines using the GSH kit. Data from three representative experiments are shown (mean SE; one-way ANOVA: *, *P* ≤ 0.05; **, *P* ≤ 0.01; ***, *P* ≤ 0.001; ****, *P* ≤ 0.0001; ns, not significant).

Next, we aimed to decipher their physiological relevance in plants by measuring total GSH levels in knockdown and overexpression lines. During the control condition, there was no substantial difference in the amount of GSH in the knockdown lines when compared with Col-0 plants [Fig. 5C]. Similarly, there was no alteration in glutathione levels in overexpression lines [Fig. 5D]. Interestingly, there was a significant reduction in the GSH pool in both the knockdown lines during oxidative stress [Fig. 5E]. On the other hand, there was a moderate increment in GSH levels in plants overexpressing *AtDJ-1C* and *AtDJ-1E* as comparison to Col-0 [Fig. 5F]. Together, these results suggest that both *At*DJ-1C and *At*DJ-1E preserve the GSH pool in yeast and plant models.

## Discussion

DJ-1 superfamily proteins are well-known for combating multiple stresses, validated in various organisms to provide cellular health and organelle protection (Lin et al., 2011a; Subedi et al., 2011; Lee et al., 2012; Richarme et al., 2018; Bankapalli et al., 2020). They are also identified in many studied organisms with varying numbers of paralogs ranging from 1 to 25, along with multiple splice variants (Ghosh et al., 2016). Interestingly, plants possess a large number of DJ-1 homologs, probably to counteract prolonged unfavourable conditions and their functions are not completely characterized. Unlike other organisms, plant DJ-1 homologs possess double DJ-1 domains tandemly arranged in a single polypeptide backbone with distinct physiological roles. This strategy may enable a stringent defence system for survival, growth and development. Our previous study revealed that double-domain *Arabidopsis* DJ-1D having two catalytic sites had enhanced functional properties (methylglyoxalase, deglycase and chaperone activity) as compared to single-domain DJ-1 proteins (Prasad et al., 2022). Since *Arabidopsis* possess multiple isoforms of the *DJ-1* gene (DJ-1A-F), they might pose scope for distinct and novel functions. Sequence alignment depicts that *At*DJ-1C lacks conserved cysteine residue in its active site, and *At*DJ-1E has only one catalytic site in its N-terminal domain (Kwon et al., 2013). In our current study, we report that *At*DJ-1C and *At*DJ-1E lack conventional methylglyoxalase and glyoxalase activity and do not attenuate the formation of MG and GO-derived advanced glycation end products (AGEs). Moreover, their expression was significantly reduced in the presence of MG stress, emphasizing the requirement of other DJ-1 paralog in (DJ-1D) under carbonyl stress.

Besides carbonyl stress, *AtDJ-1C* and *AtDJ-1E* exhibit varying expressions under different abiotic stress conditions. Furthermore, comparative transcriptome studies revealed that *AtDJ-1C* and *AtDJ-1E* were highly upregulated during heat and oxidative stress. A similar observation was also noted in different plant species, such as *Medicago truncatula*, *Glycine max*, *Erianthus arundinaceus* and *Solanum lycopersicum*, where abiotic stress influenced the expression of DJ-1 proteins (Ghosh, 2017b; Islam and Ghosh, 2018; Mohanan et al., 2020b; Gambhir et al., 2023b). Importantly, heat stress affects plant growth by altering the biochemical and physiological processes, including cell division, differentiation, respiration and photosynthesis (Giri et al., 2017; Zhao et al., 2021). Heat stress is also known to target and extensively damage chloroplasts and mitochondria; consequently, the PSII and RuBISCO activity of chloroplasts and the electron transport chain in mitochondria are being affected(Perdomo et al., 2017; Fortunato et al., 2023). Together, these global alterations contribute to the production of ROS, which damages the cellular biomolecules such as nucleic acids, proteins, lipids and pigments, leading to chronic tissue injury and cell death. Interestingly, we show that *At*DJ-1C, a chloroplast-targeted protein which is involved in chloroplast development (Lin et al., 2011b), is a robust regulator of plant health under heat stress. Its absence leads to poor root elongation, while the overexpression preserved root integrity when exposed to prolonged thermal stress. Although there could be multiple pathways to protect root length, our results strongly suggest the role of ROS scavenging as a primary defence mechanism in plant development. On the other hand, our report infers a similar method of governing plant physiology by *At*DJ-1E under heat stress, which regulates root health and robust clearance of ROS from leaves. In our studies, we show that plant DJ-1 proteins provide thermal tolerance against heat stress, while multiple studies have shown the role of DJ-1 proteins in salinity, osmotic and drought stress.

Accumulation of ROS is the primary hallmark of multiple abiotic stress, which induces metabolic dysfunction, disrupts cellular homeostasis and stunts plant development (Sachdev et al., 2021). The overexpression of *Arabidopsis* DJ-1 homologs significantly improved various health parameters of plants, such as enhanced length of plant root, enriched chlorophyll content, and suppressed ROS in leaves when exposed to toxic levels of H_2_O_2_. Interestingly, overexpression of *At*DJ-1C under control conditions retained higher chlorophyll content than Col-0 (wild-type) and DJ-1E lines, possibly protecting chloroplast through translocation and stabilizing essential proteins. On the other hand, DJ-1E acts as an antioxidant due to the presence of oxidizing cysteine (C122) in its catalytic site. Moreover, DJ-1C and DJ-1E function similarly in *S. cerevisiae*, where they provided tolerance to oxidative stress and rescued the strains lacking yeast DJ-1 member (Hsp31). Mechanistically, they maintain redox homeostasis by re-establishing the glutathione levels, a marker for oxidative stress; this finding was consistently observed in both yeast and plant models. By preserving glutathione, cells can modulate metabolic networks that may aid in countering various abiotic stress conditions. Importantly, the homozygous knockout lines of *At*DJ-1C and *At*DJ-1E could not survive, probably due to the perturbation of GSH levels that are associated with embryonic lethality (Cairns et al., 2006). Our validation in the dual model system suggests that DJ-1 members could work universally with overlapping functions despite having sequence diversity.

In conclusion, our study further sheds light on the functional diversity of plant DJ-1 proteins by exposing them to harsh environmental conditions that plants encounter in different stages of development. In particular, *At*DJ-1C and *At*DJ-1E offered enhanced rosette formation and increased plant height, suggesting that DJ-1 paralogs are interesting candidates for genetic engineering and crop improvement. Further studies can be performed exploring the role of DJ-1 paralogs in biotic stress, one of the leading factors that culminate in poor crop yield.

## Materials and methods

### Plant materials, Growth conditions and Plant Transformation

Seeds of T-DNA insertion lines *AtDJ-1C* (SALK_125439.41.70.x) and AtDJ-1E (Wiscseq_DsLox481-484P16.0) mutant lines of *Arabidopsis thaliana* (ecotype Col-0) were obtained from the Arabidopsis Biological Resource Center at Ohio State University. Surface sterilization of seeds was performed by incubating the seeds in a solution containing 0.05% SDS in 70% ethanol for 15 min and cold-stratified for 3 days and then germinated on solidified half-strength Murashige and Skoog (½ MS) media. Growth chambers (Percival) were maintained at 22 °C with long day (LD) conditions (16 h:8 h; light: dark cycle) with light intensity 100 μmol/m2/s. The germinated seedlings were transferred onto soil (composition perlite and solite in the ratio of 531 1:2). The growth room condition was maintained at 22 °C with 70% RH and long-day (LD) conditions (16 h:8 h; light: dark cycle) with light intensity 100 μmol/m2/s.

### Yeast strains and plasmid constructs

The haploid yeast BY4741 (*MATa his3Δ1 leu2Δ0 met15Δ0 ura3Δ0*) used was obtained from Open Biosystems. Haploid Δ*hsp31* and Δ*glo1* were generated by homologous recombination as previously described (Bankapalli et al., 2015; Susarla et al., 2023). *Arabidopsis thaliana DJ-1C* (without N-terminal signal sequence) and *DJ-1E* were cloned into pRS415 (Addgene) using BamHI and SalI sites under GPD (Glyceraldehyde -3-phosphate dehydrogenase) promoter with N-terminal HA-tag by using primers P1-P4 (Table 1). *AtDJ-1C* (without N-terminal signal sequence) and *AtDJ-1E* were cloned into a pRSF-duet vector (Addgene) for protein purification by using primers P5-P8 (Table 1).

For plant transformations, *AtDJ-1C* and *AtDJ-1E* were cloned in the pGWB652 with an N-terminal GFP tag through gateway cloning by using primers P12-P15 (Table 1).

### Phenotypic Analysis

Yeast strains were grown overnight in synthetic dropout media harvested at mid-log phase (*A_600_* ∼ 0.6). For methylglyoxal and glyoxal stress, cells were pelleted down at the mid-log phase and spotted on synthetic dropout media with or without 0.8 mM methylglyoxal and 15 mM glyoxal. Plates were incubated at 30 °C, and images were taken at 36 h. For oxidative stress, cells were pelleted down at the mid-log phase and resuspended in sterile water or treated with 1 mM H_2_O_2_ followed by incubation at 30 °C for 2 h. Subsequently, cells were washed with sterile water, serially diluted with a 10-fold difference and spotted on synthetic dropout media.

### *In vitro* methylglyoxalase and glyoxalase activity

To estimate the methylglyoxalase and glyoxalase activity, 5 µg of purified *At*DJ-1C, *At*DJ-1D and *At*DJ-1E were incubated with 0.5 mM MG or GO mixture containing 20 mM HEPES sodium salt (pH 7.5), 80 mM KCl, and 10% glycerol, for 45 min at 30 °C. Later, the reaction was terminated using 0.1% DNPH (2,4-dinitrophenylhydrazone), followed by adding 10% NaOH and incubating at 42 °C for 10 min. The results were calorimetrically analyzed using a spectrophotometer, 570 nm for GO and 530 nm for MG. BSA was used as the negative control.

### Measurement of ROS levels

Yeast cells were harvested from the early exponential growth phase (*A_600_* ∼ 0.5) by centrifugation and treated with 1 mM H_2_O_2_ for 1 h at 30 °C. Subsequently, the cells were treated with 100 µM H_2_DCFDA (peroxide-specific) and 10 µM DHE (superoxide-specific) for 20 min followed by a single wash with 1X PBS. To measure ROS levels by flow cytometry, treated cells were analyzed using a BD FACS cytoflex flow cytometer. The cells were subjected to fluorescence microscopy (Delta Vision Elite fluorescence microscope) for imaging.

### Measurement of Glutathione Levels

Yeast cells were grown till mid-log phase (*A_600_* ∼ 0.5) at the permissible temperature, harvested and treated with 1XPBS containing 30 µM monochlorobimane dye for 30 min at 30 °C allowing the formation of a GSH-specific adduct. Fluorescence due to adduct formation (350/460 nm, excitation/emission) was measured by flow cytometry and fluorescence microscopy (Delta Vision Elite fluorescence microscope).

Plant total glutathione levels were calculated using a glutathione GSH/GSSG calorimetric assay kit (Sigma) in control or 5 mM H_2_O_2_ treatment conditions.

### RT-PCR analysis

RNA isolation was done as previously described with minor modification (Pullagurla et al., 2023). 9-day-old seedlings were harvested in liquid nitrogen for total RNA extraction using TRIzol reagent. The seedlings were crushed with a pestle, and 1 mL of TRIzol reagent was added to homogenize them. The homogenate was incubated at room temperature for 5 min and chloroform (200 μL/mL) was added and shaken vigorously, followed by incubation of 2 min at room temperature. The homogenate was spun down at 12,000× *g* for 15 min at 4 °C. The separated aqueous phase was separated out in a new tube. To precipitate RNA, 250 μL of 100% isopropanol was added to the aqueous phase, incubated at room temperature for 10 min, and then centrifuged at 10,000× *g* for 10 min at 4 °C. The supernatant was carefully removed from the tube, without disturbing the RNA pellet. The RNA pellet was washed with 75% (*v*/*v*) ethanol and centrifuged at 7000× *g* for 5 min at 4 °C. The supernatant was discarded, and the RNA pellet was air-dried for 5–10 min at room temperature.

The total RNA extracted was then subjected to DNase1 treatment to remove genomic DNA contamination. 1 µg of RNA was used to synthesize cDNA with iScript™ cDNA Synthesis Kit (Biorad), and 100 ng cDNA was used as a template for the reaction. The qPCR was performed using the SYBR Green supermix (Biorad) with CFX96 Touch Real-Time PCR Detection System (Bio-Rad Hercules, CA, USA) by using primers P16-P19 (Table 1). The relative quantitation method (ΔΔCT) was used to evaluate quantitative variation among replicates. *β-TUBULIN* was used as the reference gene by using primers P20-P21 (Table 1).

### Western Blotting

Respective strains were grown till the mid-log phase (*A_600_*∼0.6) and treated with 2 mM MG and 12 mM GO for 12 h. The cell lysates were prepared by incubating with 10% Trichloro acetic acid solution at 4 °C followed by acetone washes. Subsequently, samples were vortexed with acid-washed glass beads and the samples were heated at 92 °C. An equal amount of sample was loaded in 12% SDS-PAGE and transferred to the PVDF membrane. The membrane was blocked with 5% BSA in phosphate-buffered saline (PBST-Tween 0.05%). Further, the membrane was incubated overnight with anti-MAGE (1:1500) or anti-CML (1:5000) antibodies at 4 °C. Following three washes with PBST, the membrane was incubated with the secondary antibody (1:15,000). Subsequently, the membrane was washed with PBST and exposed to luminol solutions (BIO-RAD). Equal loading of samples was confirmed through ponceau S staining of the membrane.

### Root length analysis

Seeds were sown on solidified half-strength MS media, supplemented with 1% (*w*/*v*) sucrose. After three days of stratification, the seedlings were grown vertically in the plant chamber under conditions of 8 h light (22 °C) and 16 h dark (20 °C) for another four days. Afterwards, seedlings were moved onto half-strength MS media supplemented with 100 mM NaCl or 100 mM Mannitol or 1 mM H_2_O_2_ or 0.6 mM H_2_O_2_ or 1 mM H_2_O_2_. For thermal and cold stress, the seedlings were transferred to 28 °C or 30 °C (for heat stress) or 4 °C (for cold stress) and maintained for the next 7 days(Yadav et al. 2023). The seedlings were allowed to grow vertically for a further seven days, and the primary root length was noted on the seventh day of growth. Images were taken using Bio-Rad ChemiDoc Version 2.4.0.03. The primary root length was measured and quantified using ImageJ 1.53k software.

### Estimation of chlorophyll content

The chlorophyll content of knockdown and overexpression lines was determined according to (Anie and Ar No N, 1949). 100 mg leaves were treated with 5 mM H_2_O_2_ (for knockdown lines) or 10 mM H_2_O_2_ (for overexpression lines) or water for 12 h. Further, they were ground in 80% acetone and centrifuged at 12000× *g* for 15 min. The absorbance of the extract was measured at 645 and 663 nm against the solvent (80% acetone) control.

### DAB staining

Five-week-old knockdown and overexpression lines plants were treated with 5 mM H_2_O_2_ for 3 h (for knockdown lines)/12 h (for overexpression lines), and left from the bolting stage were collected for DAB (3,3′-diaminobenzidine) staining as described (Daudi et al., 2012). H_2_O_2_ and water-treated leaves were immersed in DAB solution (1 mg/ml) and incubated in the dark for 5 h with shaking at 80 rpm. After incubation, the DAB solution was replaced with a bleaching solution (Ethanol: Acetic acid: Glycerol; 3:1:1). The leaves were boiled for 20 min in a water bath at 70 °C. The bleaching solution was replaced by a fresh solution and allowed to stand for 30 min.

### Statistical analysis

Statistical analyses were performed using GRAPHPAD PRISM 8.0. Bars represent SE and are derived from a minimum of three replicates. For significance testing by one-way ANOVA, Dunnett’s multiple comparison post-test was used. Asterisks represent the following significance values: *, *P* ≤ 0.05; **, *P* ≤ 0.01; ***, *P* ≤ 0.001; ****, *P* ≤ 0.0001.

### Accession numbers

Sequence data from this article can be found in the GenBank/EMBL data libraries under the accession numbers: *DJ-1C* (AT4G34020), *DJ-1D* (AT3G02720), *DJ-1E* (AT2G38860)

## Acknowledgements

We thank the Arabidopsis Biological Resource Centre (Ohio State University) for the seeds of the mutant lines that were used in this study. We thank Prof. Elizabeth A. Craig (University of Wisconsin– Madison, Madison, WI) for gifting yeast plasmids and antibody against Ydj-1.

## Supplemental Data

The following supplemental materials are available.

**Supplementary Figure S1.** Multiple Sequence alignment of *Arabidopsis* DJ-1 paralogs.

**Supplementary Figure S2.** Confirmation of knockdown lines.

**Supplementary Figure S3.** Generation of overexpression lines.

**Supplementary Table 1**

## Author Contributions

P.K. and P.D.S. perceived and designed the project. P.K. and N.J.P. performed the experiments and analyzed the data. P.D.S. and D.L. supervised the project. P.K. drafted the manuscript with inputs and approval from N.J.P., D.L. and P.D.S.

